# Sulfoquinovosylacylpropanediol monotherapy suppresses canine hemangiosarcoma patient-derived xenograft models with vascular remodeling

**DOI:** 10.64898/2026.07.03.735423

**Authors:** Keisuke Aoshima, Natsuki Miyazaki, Takuma Goto, Kazuki Heishima

## Abstract

Canine hemangiosarcoma (HSA) is an aggressive endothelial malignancy with limited therapeutic options, and its progression is closely associated with vascular architecture, stromal remodeling, and inflammatory cell recruitment. Sulfoquinovosylacylpropanediol (SQAP) is a sulfoquinovosyl lipid radiosensitizer reported to affect angiogenic and tumor-microenvironmental pathways, but its effects in canine HSA are unknown. Here, we evaluated SQAP in canine HSA cell lines and patient-derived xenograft (PDX) models. SQAP showed minimal direct cytotoxicity against HSA cell lines *in vitro*, whereas it significantly suppressed tumor growth in three canine HSA PDX models. Transcriptome analysis of SQAP-treated HSA PDX tumors detected more SQAP-responsive genes in mouse host-derived cells than in canine tumor cells. Gene-set enrichment analysis of the mouse host-derived fraction showed positive enrichment of angiogenesis, hypoxia, and stromal remodeling-related gene sets after SQAP treatment. Subsequent tissue analysis showed that SQAP reduced host-derived CD31-positive vascular area and increased α-smooth muscle actin coverage of remaining vessels in two of the three PDX models, while altering macrophage-associated marker profiles in a model-dependent manner. These findings indicate that SQAP suppresses canine HSA PDX growth primarily through vascular and macrophage-associated remodeling of the tumor microenvironment rather than direct tumor-cell cytotoxicity.

## Introduction

Canine hemangiosarcoma (HSA) is an aggressive malignant endothelial tumor with limited therapeutic options [1,2]. Although surgery and chemotherapy remain the standard treatment approaches, HSA frequently metastasizes or disseminates within body cavities, making durable disease control difficult. The limited efficacy of conventional cytotoxic agents highlights the need for additional therapeutic strategies that act through distinct targets or mechanisms of action [3–5].

The tumor microenvironment is particularly relevant to HSA because the neoplastic cells are of endothelial lineage and form blood-filled vascular structures rather than growing exclusively as solid tumor masses [2]. Thus, HSA progression is closely connected to vascular architecture, extracellular matrix, and inflammatory cell recruitment. Previous gene-expression studies support this concept, showing that inflammation and angiogenesis are distinguishing biological features of canine HSA [6]. In addition, IL-8-high HSA tumors exhibit a reactive microenvironment signature involving coagulation, inflammation, and fibrosis networks [7]. In the same study, IL-8 blockade inhibited tumor-cell engraftment *in vivo* despite limited effects on HSA cell proliferation *in vitro*, suggesting that HSA-promoting cytokine pathways may act partly through the surrounding microenvironment [7]. Furthermore, canine HSA cells have been shown to generate a stromal-immune niche that supports host-derived hematopoietic expansion [8]. Macrophages also constitute a major immune-cell population in the canine HSA microenvironment, and HSA cells can recruit macrophages and induce M2-like and PD-L1-expressing phenotypes [9]. Naturally occurring canine visceral HSAs are also enriched with tumor-associated macrophages showing an M2-like phenotype, supporting the biological relevance of macrophage-associated remodeling in this disease [10]. These observations are consistent with broader concepts in human oncology, where vascular and immune normalization of the tumor microenvironment have emerged as therapeutic strategies to improve cancer treatment [11]. This evidence indicates that the tumor microenvironment may help sustain HSA growth and could be therapeutically targeted in this disease.

Sulfoquinovosylacylpropanediol (SQAP) is a sulfoquinovosyl lipid derivative developed as an anti-cancer radiosensitizing agent. Previous studies have shown that SQAP binds focal adhesion kinase (FAK), suppresses VEGF- or fibronectin-associated FAK phosphorylation and downstream paxillin/Akt signaling, and inhibits endothelial-cell migration [12]. Beyond direct FAK signaling, SQAP has been reported to affect tumor vascular and hypoxia-related pathways in xenograft models. SQAP monotherapy suppressed angiogenesis and hypoxia-related signaling in hepatoma xenografts through pVHL upregulation and HIFα downregulation [13]. In radiotherapy-combination models, SQAP enhanced radiation response and was associated with vascular or hypoxia-related remodeling in prostate cancer and malignant mesothelioma xenografts [14,15]. Direct functional evidence for SQAP-induced vascular effects comes from a multimodal imaging study showing acute increases in tumor perfusion and pOL after SQAP administration and reduced microvascular density after repeated SQAP-based treatment [16]. More recent work further suggests that SQAP can interfere with DNA repair pathways and topoisomerase activity, thereby enhancing radio- and chemosensitivity in canine cancer cell lines [17]. These studies suggest that SQAP may act through both direct effects on tumor cells and modulation of the tumor microenvironment. However, the effects of SQAP in canine HSA have not been examined. Since HSA is an endothelial malignancy that forms neoplastic vascular structures, SQAP could plausibly affect HSA by acting directly on neoplastic endothelial cells, by modifying host-derived vascular compartments, or by both mechanisms.

In the present study, we evaluated the effects of SQAP on canine HSA cell lines and canine HSA patient-derived xenograft (PDX) models. We previously established a canine HSA PDX platform from surgically resected patient tumors and showed that these models retained the histological features of the corresponding original tumors [18,19]. Since these PDX models maintain patient-derived tumor architecture in an *in vivo* setting, they provide a pathologically relevant platform for evaluating the effects of SQAP on canine HSA.

## Results

### SQAP suppressed canine hemangiosarcoma PDX tumor growth despite minimal direct cytotoxicity in vitro

To examine whether SQAP directly affects canine hemangiosarcoma cell viability, HU-HSA-2 and HU-HSA-3 cells were treated with SQAP or doxorubicin for 48 h. SQAP did not affect cell viability in either cell line, whereas doxorubicin, used as a cytotoxic control, markedly reduced cell viability in both cell lines (Fig. 1A). In contrast, SQAP significantly suppressed tumor growth in HSA PDX models (Fig. 1B, D, F), and no difference was observed between the 4 mg/kg and 12 mg/kg SQAP treatment groups. Analysis of mean tumor volume further supported the anti-tumor effect of SQAP (Fig. 1C, E, G). Using phosphate-buffered saline (PBS)-treated tumors as the control reference, most PDX tumors treated with SQAP were classified as responders, although some were classified as worse-than-PBS or PBS-like non-responders. This finding may indicate model- and tumor-level heterogeneity in treatment response. Representative hematoxylin and eosin (H&E)-stained sections of major organs collected at necropsy did not reveal overt treatment-associated histopathological alterations in SQAP-treated mice under the examined conditions (Supplementary Figs. S1–S3).

**Figure 1.**
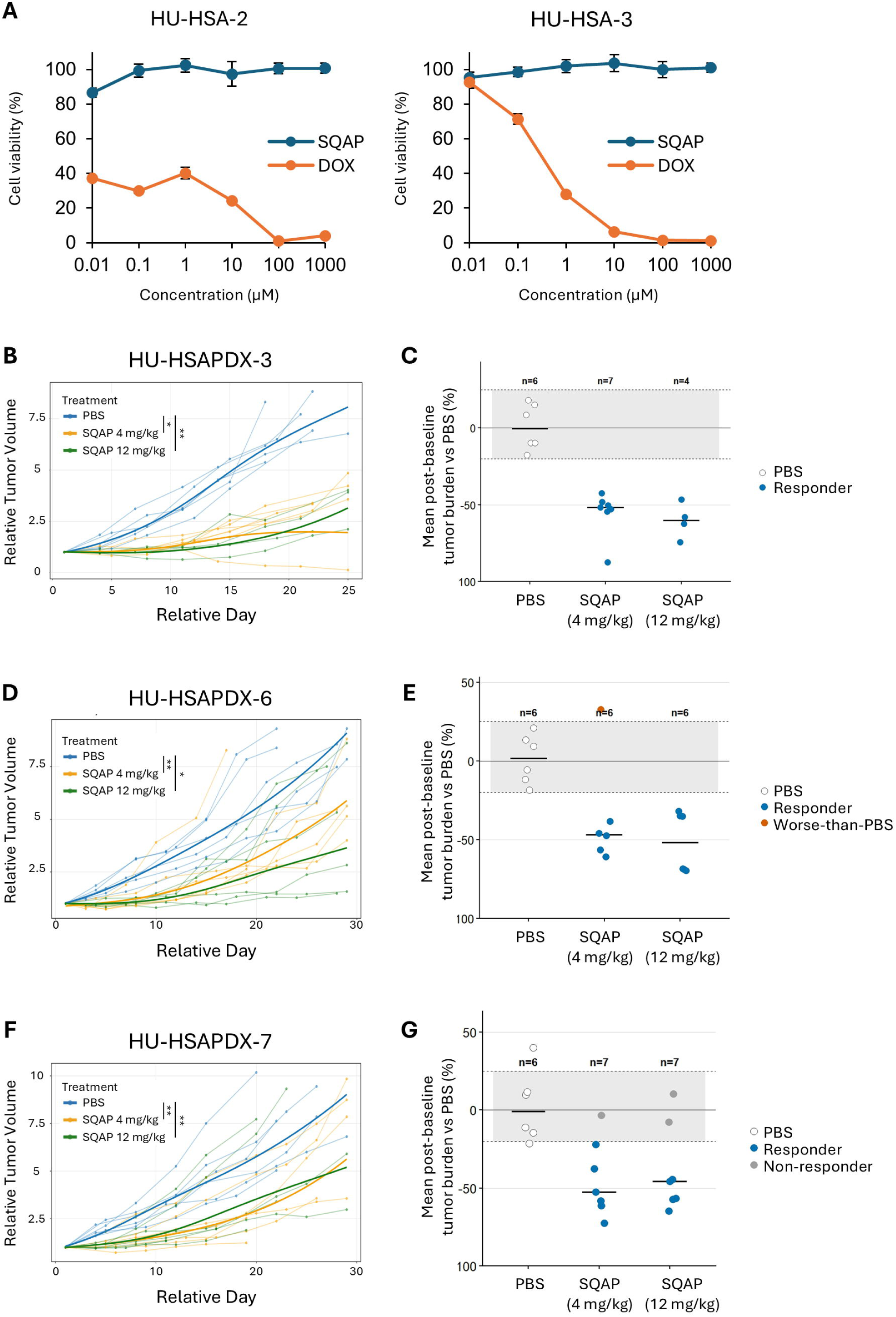
SQAP suppresses tumor growth in canine HSA PDX models but not in cell lines. **(A)** HU-HSA-2 and HU-HSA-3 cells were treated with SQAP or doxorubicin (DOX) with indicated concentrations for 48 h. Viability was normalized to untreated control cells after background correction. Data are shown as mean ± SD from triplicate wells. **(B, D, F)** Growth curves of HU-HSAPDX-3, HU-HSAPDX-6, and HU-HSAPDX-7 tumors in KSN/Slc mice treated with PBS, SQAP 4 mg/kg, or SQAP 12 mg/kg. Tumor volume was normalized to the volume at treatment initiation (day 1 = 1). Thick lines show smoothed model-based estimates of the mean relative tumor volume. Thin lines indicate individual tumor volumes. * *p* < 0.05; ** *p* < 0.01. Adjusted *p* values for SQAP 4 mg/kg and 12 mg/kg versus PBS were 0.0007 and 0.0016 for HU-HSAPDX-3, 0.0102 and 0.0006 for HU-HSAPDX-6, and 0.0034 and 0.0168 for HU-HSAPDX-7, respectively. **(C, E, G)** Mean tumor volume relative to PBS controls. Each point represents one mouse. Values were calculated by comparing each tumor volume measurement with the PBS control value and averaging across measurement periods. The shaded area indicates the predefined control range (0.80 to 1.25 times the PBS control level). Treated mice below this range were classified as responders, mice within this range as non-responders, and mice above this range as worse than control.

Next, we performed messenger RNA sequencing (mRNA-seq) of SQAP-treated canine hemangiosarcoma cell lines and HU-HSAPDX-6 tumors to examine the effects of SQAP on gene expression. Given that SQAP rapidly disappears from plasma but is relatively retained in tumor tissues [20], two time points were evaluated to analyze acute and delayed exposure effects: 30 min and 3 days for *in vitro* experiments, and 60 min and 3 days for *in vivo* experiments. In PDX tumor samples, canine tumor-derived reads and mouse host-derived reads were separated and analyzed independently to evaluate the effects of SQAP on tumor cells and non-tumor cells, respectively. Under *in vitro* conditions, 30-min SQAP treatment produced minimal changes in gene expression in both cell lines (Fig. 2A and C). Three-day SQAP treatment altered gene expression in HU-HSA-2 (80 upregulated genes and 34 downregulated genes); however, this response was not consistently observed in HU-HSA-3, which showed a limited response (no upregulated genes and 3 downregulated genes) (Fig. 2B and D). In PDX tumors, SQAP produced relatively limited differential gene expression in the canine tumor-cell fraction. At 60 min, no upregulated genes and 6 downregulated genes were identified; after 3 days, 2 upregulated genes and 5 downregulated genes were identified (Fig. 2E and F). By contrast, the mouse host-cell fraction showed broader transcriptional alterations. At 60 min, 9 upregulated genes and 37 downregulated genes were identified, and after 3 days, 23 upregulated genes and 32 downregulated genes were identified (Fig. 2G and H). These data indicate that the *in vivo* transcriptional response to SQAP involved more SQAP-responsive genes in mouse host-derived cells than in canine tumor cells.

**Figure 2.**
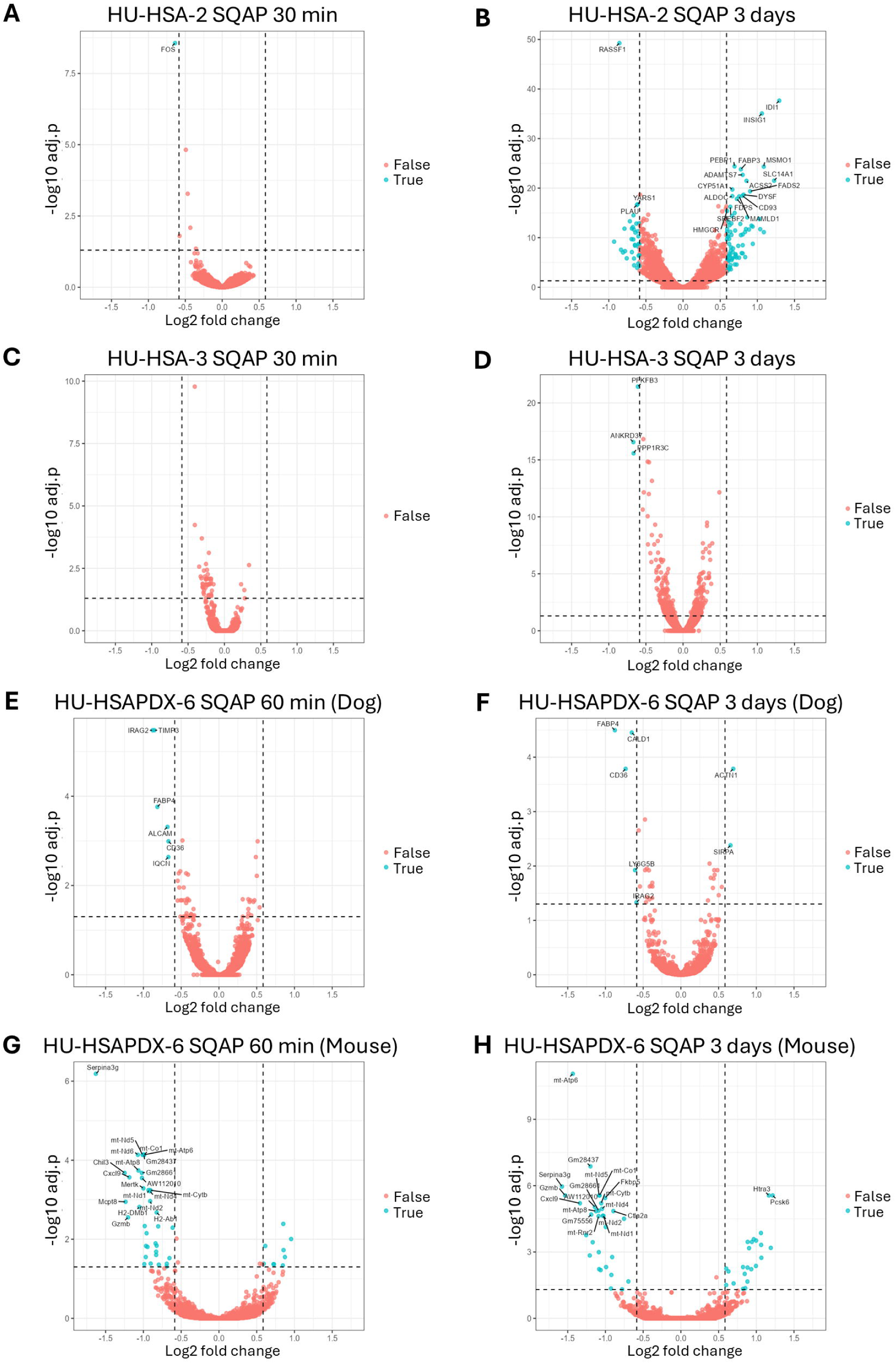
Acute and delayed gene expression changes to SQAP in canine hemangiosarcoma cells and HU-HSAPDX-6 tumors. Volcano plots showing differential gene expression after SQAP treatment in vitro and in vivo. **(A, B)** HU-HSA-2 cells treated with SQAP for 30 min or 3 days. **(C, D)** HU-HSA-3 cells treated with SQAP for 30 min or 3 days. **(E, F)** Canine tumor-cell component of HU-HSAPDX-6 tumors after SQAP treatment for 60 min or 3 days. **(G, H)** Mouse host-cell component of the same HU-HSAPDX-6 tumors after SQAP treatment for 60 min or 3 days. Each condition was analyzed in triplicate. Dashed vertical and horizontal lines indicate |log□FC| ≥ 0.585 and FDR-adjusted *p* < 0.05, respectively. Cyan points indicate genes meeting both significance thresholds.

These results suggest that SQAP suppresses HSA PDX tumor growth largely through modulation of the tumor microenvironment rather than through overt direct cytotoxicity to tumor cells.

### SQAP remodels the PDX tumor microenvironment

Given that SQAP treatment reduced PDX tumor growth and markedly affected gene expression profiles in mouse-derived host cells, we performed gene-set enrichment analysis (GSEA) to identify the pathways modulated by SQAP. In the mouse host-cell fraction of HU-HSAPDX-6 tumors following the 3-day SQAP treatment, GSEA showed positive enrichment of gene sets associated with tumor microenvironment remodeling, including angiogenesis, hypoxia, and epithelial-mesenchymal transition (EMT)-associated stromal remodeling (Fig. 3A). These pathway-level changes suggested that SQAP altered vascular- and stromal-associated programs in the host tumor microenvironment. Since previous SQAP xenograft studies have linked SQAP treatment to angiogenic, hypoxia-related, and vascular-remodeling phenotypes [13–16], we next examined whether SQAP treatment was associated with structural remodeling of host-derived tumor vasculature in endpoint PDX tissues. To this end, PDX tumor tissues from the efficacy study shown in Figure 1 were stained for the endothelial marker CD31 using clone D8V9E, which recognizes murine but not canine CD31, and for α-smooth muscle actin (αSMA) to assess host-derived vascular area and mural-cell coverage. Quantification of CD31 signals in viable tumor regions indicated that SQAP reduced CD31-positive area density in HU-HSAPDX-3 and HU-HSAPDX-7 tumors compared with PBS controls, although HU-HSAPDX-6 did not show a reduction. We further measured αSMA coverage in tumor blood vessels to evaluate vascular maturity. SQAP significantly increased αSMA coverage of CD31-positive vessels in HU-HSAPDX-3 and HU-HSAPDX-7 tumors compared with PBS controls (Fig. 3C, D). In contrast, αSMA coverage was not significantly altered in HU-HSAPDX-6 tumors. Thus, increased αSMA coverage was observed in the two PDX models in which SQAP reduced CD31-positive vascular area.

**Figure 3.**
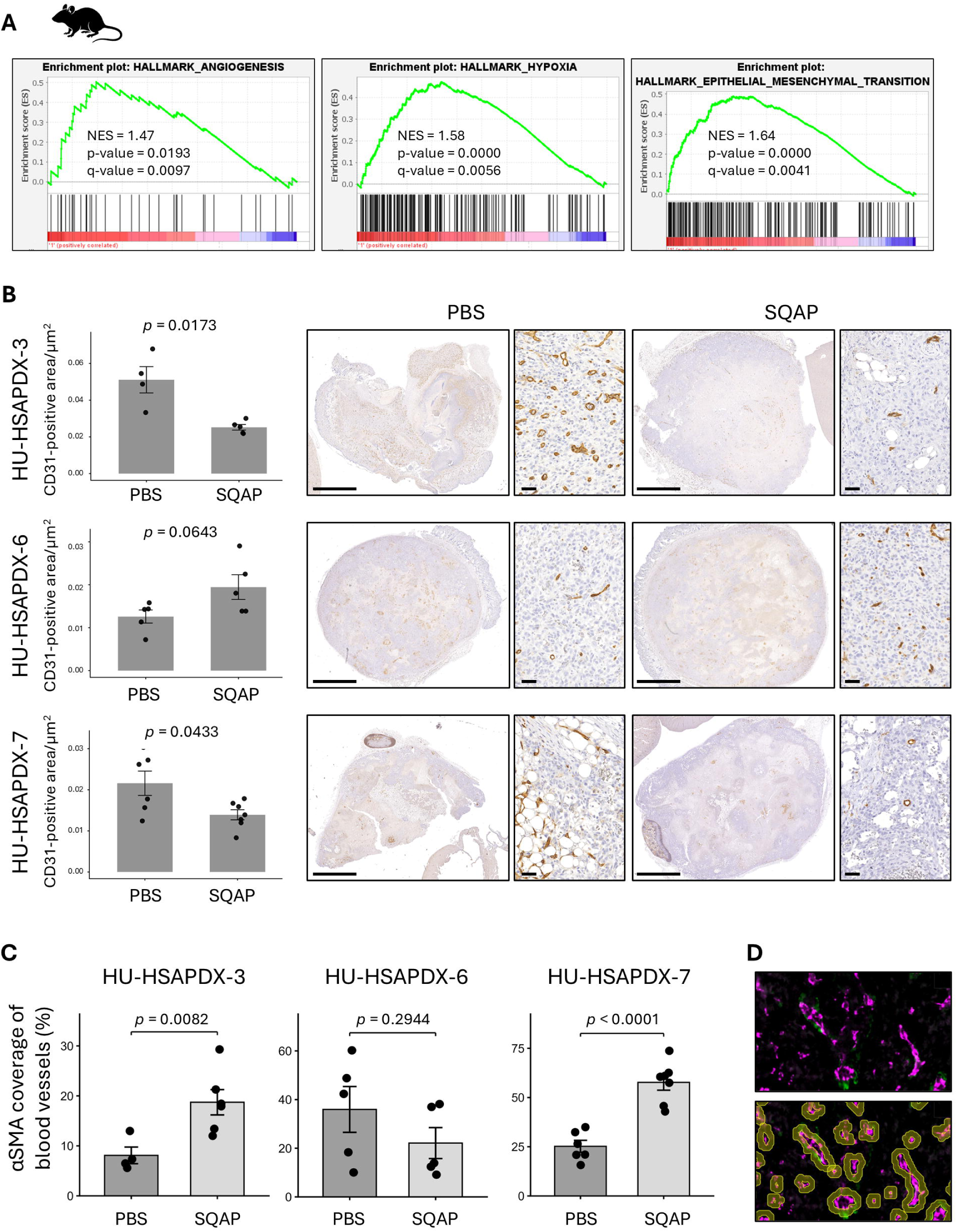
SQAP alters host vascular density and αSMA coverage in canine hemangiosarcoma PDX tumors. **(A)** Gene set enrichment analysis of the mouse host-derived gene expressions from HU-HSAPDX-6 tumors comparing 3-day SQAP-treated tumors with PBS controls. NES, normalized enrichment score. **(B)** (Left) Quantification of CD31-positive area in viable tumor regions of tumor tissues. Dots represent individual tumors, and bars show group mean ± SEM. CD31-positive area density was compared using two-sided Welch’s *t*-tests after log□□ transformation. (Right) Representative CD31 IHC images from PBS- and SQAP-treated tumors are also shown. Scale bars indicate 2.5 mm for whole-section images and 50 μm for high-magnification images. **(C)** Quantification of αSMA coverage of CD31-positive blood vessels in viable regions of tumor tissues. Dots represent individual tumors, and bars show group mean ± SEM. αSMA coverage was compared using two-sided Welch’s *t*-tests after logit transformation. **(D)** Representative CD31/αSMA dual immunofluorescence image (upper) and corresponding image-analysis overlay showing segmented CD31-positive vessel profiles and vessel-wall regions used to quantify αSMA coverage (lower). CD31 is shown in magenta, αSMA in green and DAPI in blue.

GSEA of the canine tumor-cell fraction showed that SQAP treatment was associated with negative enrichment of interferon-response pathways (Fig. 4A), which suggests that SQAP may alter interferon-related signaling in HSA tumor cells. Since macrophages constitute a major immune-cell population in the canine HSA microenvironment, macrophage-associated components of endpoint PDX tissues were then evaluated by immunohistochemistry. PDX tumor sections were stained for Iba1 and F4/80 as pan-macrophage markers, CD86 as an M1-like macrophage marker, and CD206 as an M2-like macrophage marker. The results indicated that SQAP altered macrophage marker-positive area density in a model-dependent manner (Fig. 4B). In HU-HSAPDX-3, SQAP significantly reduced Iba1-, F4/80-, and CD206-positive area densities, whereas CD86-positive area density was not altered. In HU-HSAPDX-6, Iba1-, F4/80-, and CD206-positive area densities were not significantly altered. CD86-positive area density was higher in the SQAP-treated group, but the difference did not reach statistical significance (*p* = 0.062). In HU-HSAPDX-7, Iba1- and CD206-positive area densities were not significantly changed, whereas F4/80- and CD86-positive area densities were increased after SQAP treatment.

**Figure 4.**
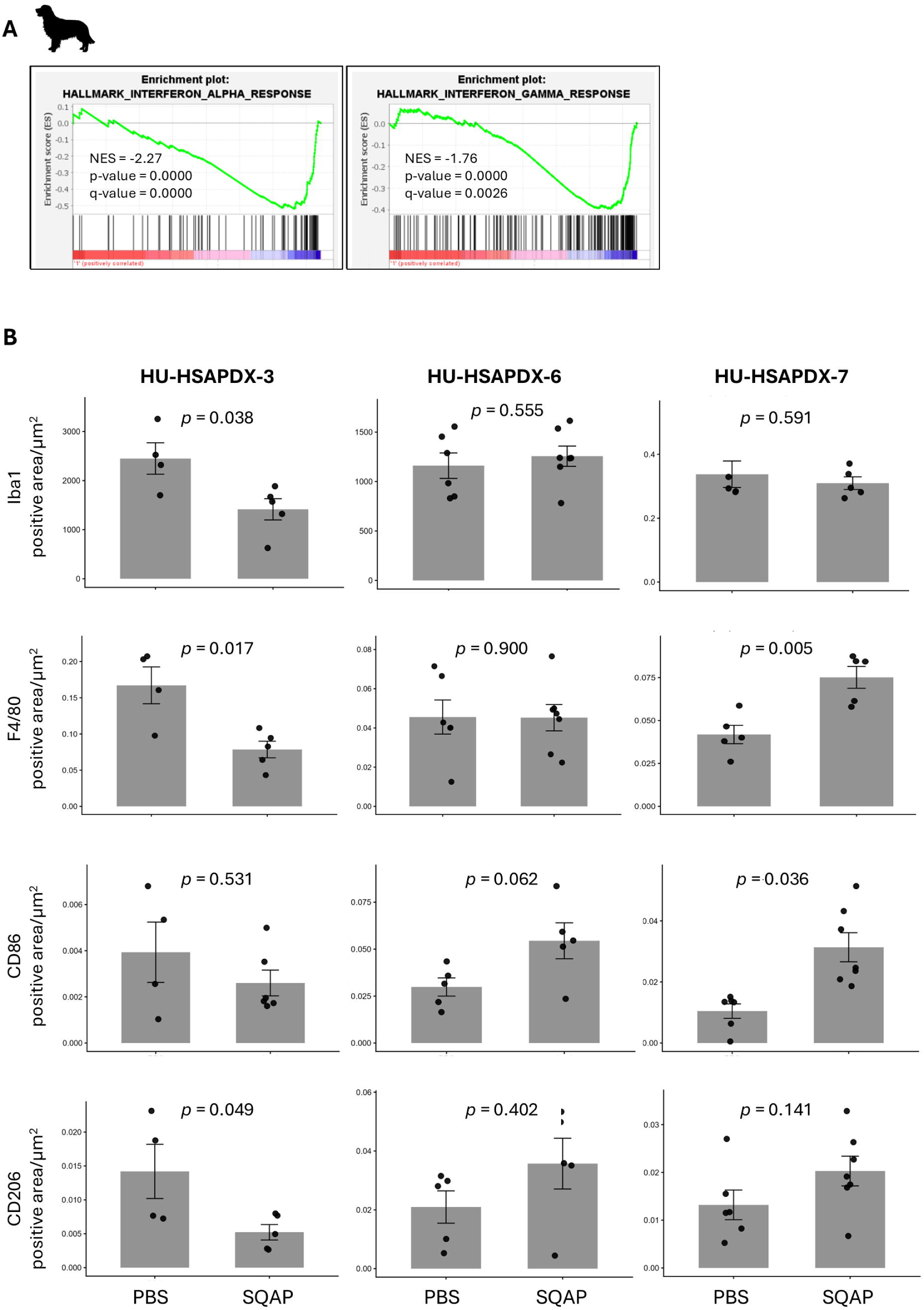
SQAP alters macrophage-associated marker profiles in PDX tumors. **(A)** Gene set enrichment analysis of the canine tumor-derived gene expressions from HU-HSAPDX-6 tumors comparing 3-day SQAP-treated tumors with PBS controls. NES, normalized enrichment score. **(B)** Quantification of macrophage-associated markers in viable tumor regions of endpoint tumor tissues collected from the efficacy study shown in Figure 1. Dots represent individual tumors, and bars show group mean ± SEM. Marker-positive area density was compared between PBS and SQAP 4 mg/kg groups within each PDX model using two-sided Welch’s *t*-tests after log□□ transformation.

Together, these results indicate that SQAP remodels both vascular and macrophage-associated components of the canine hemangiosarcoma PDX tumor microenvironment. However, the direction and magnitude of these changes varied across PDX models, suggesting that SQAP-induced microenvironmental remodeling is context dependent.

## Discussion

In this study, we found that SQAP inhibited the growth of canine HSA PDX tumors despite showing minimal direct cytotoxicity toward HSA cell lines *in vitro*. This discrepancy suggests that the anti-tumor activity of SQAP in HSA is unlikely to be explained primarily by direct cytotoxicity under the conditions tested. The reduced host CD31-positive area and increased αSMA coverage observed in HU-HSAPDX-3 and HU-HSAPDX-7 are compatible with prior SQAP studies reporting vascular and hypoxia-associated effects in xenograft models [13–16]. Importantly, the present findings extend these observations to SQAP monotherapy in canine HSA PDX models, but they should be interpreted as structural vascular remodeling rather than evidence of functional vascular normalization. Since αSMA coverage was measured only among the remaining CD31-positive vessels, the increase could reflect recruitment or stabilization of αSMA-positive mural cells, selective loss of αSMA-poor immature vessels, or both. Direct assessment of vessel perfusion, vascular leakage, and tumor hypoxia will be required to determine whether these structural changes improve blood flow or oxygenation. Unlike HU-HSAPDX-3 and HU-HSAPDX-7, HU-HSAPDX-6 did not show significant changes in CD31-positive vascular area or αSMA coverage. Since tissues for vascular analysis were collected only at the end of the efficacy study, this result does not exclude an earlier or transient vascular response. The early transcriptomic changes detected in the mouse host-cell fraction of HU-HSAPDX-6 indicate that SQAP affected its host microenvironment, but endpoint CD31/αSMA analysis may have missed dynamic vascular changes or subsequent stromal adaptation. Mechanisms not captured by CD31 and αSMA measurements may also contribute to SQAP activity in this model. Recent work showed that SQAP inhibited non-homologous end joining and homologous recombination in reporter assays, disrupted topoisomerase I and IIα activities, and enhanced radio- and chemosensitivity in canine cancer cell lines [17]. Given that HU-HSA-2 and HU-HSA-3 cells may not fully recapitulate the tumor-intrinsic biology of HU-HSAPDX-6, direct effects of SQAP on HU-HSAPDX-6 tumor cells cannot be excluded.

In parallel with vascular remodeling, SQAP altered macrophage-associated marker profiles in a model-dependent manner. Macrophage marker-positive areas decreased in HU-HSAPDX-3, F4/80- and CD86-positive areas increased in HU-HSAPDX-7, and no statistically significant changes were detected in HU-HSAPDX-6, although CD86 showed an upward trend. Given the established relevance of macrophages in canine HSA [8–10], these findings suggest that SQAP may affect macrophages either directly or secondarily to vascular remodeling. However, since we used athymic nude mice and only one marker each for M1- and M2-like phenotypes, these data do not define macrophage polarization or its functional consequences. Studies in immunocompetent settings, including tumor tissues from SQAP-treated dogs with spontaneous HSA and appropriate syngeneic mouse models, will be needed to clarify how SQAP affects macrophage biology in this disease.

Overall, these findings support a model in which SQAP suppresses canine HSA PDX growth mainly through tumor-microenvironmental effects rather than direct tumor-cell cytotoxicity. Further studies using time-course sampling, functional vascular assays, and immunocompetent models are needed to define how SQAP remodels the HSA microenvironment and to identify biomarkers associated with treatment response.

## Materials and Methods

### Cell culture and reagent preparation

Canine hemangiosarcoma cell lines HU-HSA-2 and HU-HSA-3 were cultured in high-glucose Dulbecco’s Modified Eagle Medium (DMEM; FUJIFILM Wako Pure Chemical Corporation, Osaka, Japan; Cat. No. 044-29765) supplemented with 10% fetal bovine serum (FBS; Thermo Fisher Scientific, Waltham, MA, USA; Cat. No. 10270-106) and penicillin-streptomycin solution (FUJIFILM Wako Pure Chemical Corporation; Cat. No. 168-23191) at 37 °C in 5% CO□. SQAP (Lavurchin injection 40 mg; Malignant Tumor Treatment Technologies, Inc. [M.T.3], Sakai, Osaka, Japan) was provided by ASCO Co., Ltd. (Tokyo, Japan), dissolved in sterile PBS, and serially diluted in complete DMEM immediately before treatment. Doxorubicin hydrochloride (FUJIFILM Wako Pure Chemical Corporation; Cat. No. 040-21521) was prepared as a 10 mg/mL stock solution and serially diluted in complete DMEM immediately before treatment.

### Cell viability assay

HU-HSA-2 and HU-HSA-3 cells were seeded into 96-well cell culture plates (Greiner Bio-One, Kremsmünster, Austria; Cat. No. 655180) at densities of 3.0 × 10³ cells/well and 1.5 × 10³ cells/well, respectively, and were allowed to attach overnight. Twenty-four hours after seeding, the culture medium was replaced with complete DMEM containing SQAP or doxorubicin at final concentrations of 0.01, 0.1, 1, 10, 100, or 1000 µM. Untreated cell-control wells, medium-only blank wells, and cell-free doxorubicin-containing blank wells were included for background correction. Each treatment condition was assayed in triplicate wells. Cell viability was assessed using the Cell Counting Kit-8 (CCK-8; Dojindo, Kumamoto, Japan; Cat. No. 343-07623). After 48 h of drug exposure, 10 µL of CCK-8 solution was added directly to each well. Plates were incubated at 37 °C for 4 h, and absorbance was measured at 450 nm using a microplate reader (MTP-320; Corona Electric Co., Ltd., Ibaraki, Japan). Absorbance values were corrected using the corresponding blank values. For SQAP-treated wells, medium-only blanks were used for background correction. For doxorubicin-treated wells, concentration-matched cell-free doxorubicin-containing blank wells were used to correct for drug-associated background absorbance. Cell viability was calculated relative to untreated control cells as follows: cell viability (%) = [(A□□□ of treated wells − A□□□ of the corresponding blank wells)/(A□□□ of untreated control wells − A□□□ of medium-only blank wells)] × 100. Data are presented as the mean ± standard deviation of triplicate wells. Survival curves were plotted against the logarithm of the drug concentration.

### PDX treatment with SQAP

All mouse experiment protocols were approved by Hokkaido University Institutional Animal Care and Use Committee (protocol numbers: 21–0062 and 25-0062), and conducted in accordance with the Animal Research: Reporting of In Vivo Experiments (ARRIVE) guidelines [21]. Three canine hemangiosarcoma PDX models HU-HSAPDX-3, HU-HSAPDX-6, and HU-HSAPDX-7 were previously established by subcutaneous transplantation of surgically resected splenic hemangiosarcoma tissues into 4- to 8-week KSN/Slc mice (Japan SLC, Inc., Shizuoka, Japan) [18,19]. The original tumors were obtained at Hokkaido University Veterinary Teaching Hospital from an 8-year-old spayed female Flat-Coated Retriever, a 13-year-old neutered male Chihuahua, and a 10-year-old male mixed-breed dog, respectively. For PDX transplantation, tumor tissues were cut into approximately 2–3 mm³ fragments and implanted subcutaneously into the right flank of recipient KSN/Slc mice under anesthesia with medetomidine (0.3□mg/kg), midazolam (4□mg/kg), and butorphanol (5□mg/kg). A 7-mm skin incision was made on the flank, one tumor fragment was placed subcutaneously, and the incision was closed with a surgical clip. Postoperative analgesia was provided with meloxicam (0.2 mg/kg, intraperitoneally), and surgical clips were removed 1 week after transplantation. Tumor volume was calculated as: tumor volume = length × width² / 2. Mice were euthanized with CO□ when tumors reached 1 cm³ or when endpoint criteria were met.

For the treatment study, mice bearing HU-HSAPDX-3, HU-HSAPDX-6, or HU-HSAPDX-7 tumors were monitored until individual tumor volume reached 100 mm³, which was used as the threshold for treatment initiation. Once tumors reached this threshold, mice were assigned to PBS or SQAP treatment at 4 or 12 mg/kg. Tumor volume was analyzed relative to the volume at treatment initiation. The SQAP doses were selected based on prior SQAP mouse xenograft studies [13–16]. SQAP or PBS was administered intraperitoneally five times per week (5 consecutive treatment days followed by 2 days without treatment). Tumor growth was monitored by caliper measurement twice a week, and relative tumor volume was calculated by normalizing each tumor-volume value to the volume at treatment initiation.

Statistical analysis was performed using R v4.5.3 in RStudio v2026.01.2+418. Relative tumor volume was log-transformed before analysis. The primary longitudinal analysis for overall tumor growth used an additive linear mixed-effects model fitted by maximum likelihood. Day was treated as a categorical factor, and subject was included as a random intercept. Estimated marginal means for each treatment group were calculated with equal weighting across post-baseline visits. Treatment effects for SQAP-treated groups versus PBS were evaluated using Dunnett-adjusted contrasts with the multivariate *t* adjustment. The 4 mg/kg and 12 mg/kg SQAP groups were directly compared as an exploratory dose-comparison analysis. In addition, a treatment-by-day interaction model was fitted as a sensitivity analysis to determine whether the treatment effect varied over time. For visualization, individual tumor-growth trajectories were plotted, and smooth treatment-level trajectories were overlaid using a visualization-only spline mixed model. Statistical inference was based on the categorical-day mixed model.

For responder classification, day-matched PBS reference values were estimated by PBS-treated mice using a PBS-only mixed-effects model. For each treated mouse, relative tumor volumes after treatment initiation were compared with the corresponding PBS reference values, and the results were used to calculate a PBS-adjusted tumor-burden ratio and percent difference for each mouse. Treated mice with PBS-adjusted tumor burden below 80% of the PBS reference were classified as responders, those with tumor burden between 80% and 125% of the PBS reference were classified as PBS-like non-responders, and those with tumor burden above 125% of the PBS reference were classified as worse-than-PBS. The 80%–125% range corresponds to a symmetric margin of ±log(1.25) on the log-ratio scale. Fisher’s exact test was used to compare response classifications between treated groups. PBS-like non-responders were further compared with PBS controls using Welch’s two one-sided tests based on the normalized post-baseline area under the curve for log-transformed relative tumor volume. The equivalence margin was defined as a ratio of 1.25.

### mRNA-seq

SQAP-induced gene expression changes were analyzed in canine hemangiosarcoma cell lines and an HSA PDX model (HU-HSAPDX-6). Each group consisted of three biological replicates. Given that direct measurement of free SQAP concentration in mouse plasma or tumor after intraperitoneal administration was not available, exact *in vivo*–*in vitro* concentration matching was not attempted. Instead, the *in vitro* conditions were selected as pharmacodynamic exposure models. A published radiolabeled SQAP mouse study showed rapid decline of SQAP-derived radioactivity from plasma with relatively prolonged tumor-associated retention after intravenous administration [20]. Therefore, both acute and delayed exposure effects were evaluated. For cell-line experiments, the acute 30-min exposure used 12 µg/mL SQAP to represent a high-exposure state, whereas the delayed exposure used 6 µg/mL SQAP to model a lower repeated-exposure state. These concentrations were selected to remain within a non-cytotoxic range in canine hemangiosarcoma cells (Fig. 1) and to approximate the low-micromolar exposure range used in prior SQAP mechanistic studies [12,13,17]. Since SQAP is expected to be substantially protein-bound in plasma and binding in culture medium containing 10% FBS may differ, these *in vitro* exposures were interpreted as practical pharmacodynamic models rather than direct concentration matches to the *in vivo* condition. The acute 30-min *in vitro* time point and 60-min *in vivo* time point were selected to capture early transcriptional responses, whereas the 3-day treatment followed by 24 h of drug-free culture or post-dose harvest was selected to assess delayed gene expression changes after repeated exposure. For the PDX mRNA-seq experiment, SQAP was administered at 4 mg/kg, corresponding to the lower dose evaluated in the PDX efficacy study (Fig. 2).

HU-HSA-2 and HU-HSA-3 cells were seeded in 6-well plates at 2.0 × 10□ cells/well and 1.0 × 10□ cells/well, respectively, and treated with vehicle (DMEM) or SQAP. For the acute condition, cells were treated with 12 µg/mL SQAP for 30 min without washout and harvested immediately. For the delayed condition, cells were treated with 6 µg/mL SQAP for 3 days. The medium was replaced daily with freshly prepared SQAP-containing complete medium. Cells were then cultured in drug-free complete medium for 24 h before harvest. KSN/Slc nude mice bearing HU-HSAPDX-6 tumors were treated intraperitoneally with PBS or SQAP at 4 mg/kg. Three groups were prepared: PBS-treated controls, SQAP-treated 60 min after a single dose, and SQAP-treated tumors harvested 24 h after the final dose of once-daily dosing for 3 consecutive days. Mice were anesthetized using the aforementioned medetomidine–midazolam–butorphanol method, and tumor tissues were collected. For the 60-min group, tumor tissues were harvested 60 min after SQAP treatment. For the PBS control and 3-day SQAP groups, mice were treated once daily at the same clock time, and tumors were harvested 24 h after the final treatment. Tumor tissues were maintained in DMEM at 4 °C and dissociated in 3 mg/mL collagenase I solution (FUJIFILM Wako Pure Chemical Corporation; Cat. No. 035-1760) using a VIA Extractor (Cytiva, Marlborough, MA, USA) for 10 min at 37 °C. Tissue suspensions were passed through a 70-µm cell strainer and washed with PBS. Cell pellets were washed three times with red blood cell (RBC) lysis buffer containing 0.747% NH□Cl and 17 mM Tris-HCl, pH 7.5, followed by three PBS washes. Cell pellets were stored at −80 °C until RNA extraction.

Total RNA was extracted using the NucleoSpin RNA kit (Takara Bio Inc., Shiga, Japan) according to the manufacturer’s instructions. Purified RNA was stored at −80 °C until submitting to Rhelixa, Inc. (Tokyo, Japan) for library preparation and sequencing. RNA concentration was measured, and RNA quality was assessed by electrophoresis before library preparation. Poly(A)-positive mRNA was enriched using the NEBNext Poly(A) mRNA Magnetic Isolation Module (New England Biolabs, Ipswich, MA, USA), and strand-specific libraries were prepared using the NEBNext Ultra II Directional RNA Library Prep Kit (New England Biolabs). Libraries were sequenced on an Illumina NovaSeq X Plus platform (Illumina, San Diego, CA, USA) to generate 150-bp paired-end reads. Sequencing was specified to yield 6 Gb per sample, corresponding to 40 million reads per sample, or 20 million paired-end read pairs per sample.

For canine cell-line samples, paired-end reads were aligned to the canine reference genome ROS_Cfam_1.0/CanFam4. For HU-HSAPDX-6 tumor samples, a species-disambiguated workflow was used to separate canine tumor-derived reads and mouse host-derived reads. Paired-end reads from PDX samples were first aligned independently to the canine reference genome ROS_Cfam_1.0/CanFam4 and the mouse reference genome GRCm39 with GENCODE M38 annotation using STAR v2.7.11b [22]. To separate graft-derived canine reads and host-derived mouse reads, alignments were processed using Disambiguate with the STAR aligner option [23]. Species-specific BAM files were name-sorted and converted to paired FASTQ files using SAMtools [24]. Disambiguated reads were then re-aligned to the corresponding species-specific reference genome using STAR. Gene-level abundance was estimated using RSEM [25], and canine and mouse matrices were processed separately for downstream analysis. Differential-expression analysis was performed using DESeq2 [26]. For each contrast, Wald tests were used to test whether gene-level log□ fold changes differed from zero, and *p*-values were adjusted using the Benjamini–Hochberg method to control the false-discovery rate [27]. Log□ fold-change estimates were moderated using DESeq2 lfcShrink with the normal prior and were used for volcano-plot visualization. Genes were highlighted as significant when adjusted *p* < 0.05 and |log□ fold change| ≥ 0.585, corresponding to approximately 1.5-fold change. Genes were labeled by gene symbol, or by Ensembl ID when no gene symbol was available. Gene set enrichment analysis was performed for pathway-level analysis [28–31]. Raw sequencing data and processed expression matrices are deposited in the NCBI Gene Expression Omnibus under accession number GSE336925.

### Histopathological analysis

Tumor tissues and major organs were fixed in 10% neutral-buffered formalin, dehydrated through a graded ethanol series, cleared with xylene, and infiltrated with paraffin wax using a Tissue-Tek VIP5 Jr tissue processor (Sakura Finetek Japan Co., Ltd., Tokyo, Japan). Paraffin-embedded tissues were sectioned at 3 μm thickness. Sections were deparaffinized with xylene, rehydrated through graded ethanol, and washed with PBS. For H&E staining, sections were stained with hematoxylin for 1 min and washed with tap water for 5 min. Sections were then placed in 95% ethanol for 2 min, stained with eosin for 1.5 min, and washed with 95% ethanol to remove excess eosin. After staining, sections were dehydrated with absolute ethanol, cleared with xylene, mounted with Eukitt mounting medium (ORSAtec, Bobingen, Germany), and sealed with cover glasses. For immunohistochemistry (IHC), heat-induced epitope retrieval was performed in ImmunoActive Antigen Retrieval Solution pH 6.0 (IA6500; Matsunami Glass Ind., Ltd., Osaka, Japan). For CD31 staining, antigen retrieval was performed by autoclaving at 121°C for 1 min, followed by natural cooling. For macrophage-marker staining, antigen retrieval was performed using a 2100-Retriever (Aptum Biologics, Southampton, UK). Endogenous peroxidase activity was blocked with 0.3% hydrogen peroxide in methanol for 15 min, and sections were blocked with normal goat serum (Nichirei Biosciences, Tokyo, Japan) for 30 min at room temperature (RT). Sections were incubated with primary antibodies overnight at 4°C. Primary antibodies used in this study are listed in Table 1. After washing with PBS, sections for CD31 staining were incubated with ImmPRESS-HRP Horse Anti-Rabbit IgG Polymer Detection Kit, Peroxidase (MP-7401-50; Vector Laboratories, Newark, CA, USA) for 20 min at RT. Sections stained for macrophage-associated markers were incubated with Simple Stain MAX-PO (R) (Nichirei Biosciences) for 30 min at RT. Signals were visualized with 3,3′-diaminobenzidine (DAB). Sections were then counterstained with hematoxylin, dehydrated, cleared with xylene, mounted with Eukitt mounting medium, and sealed with cover glasses.

**Table 1.**
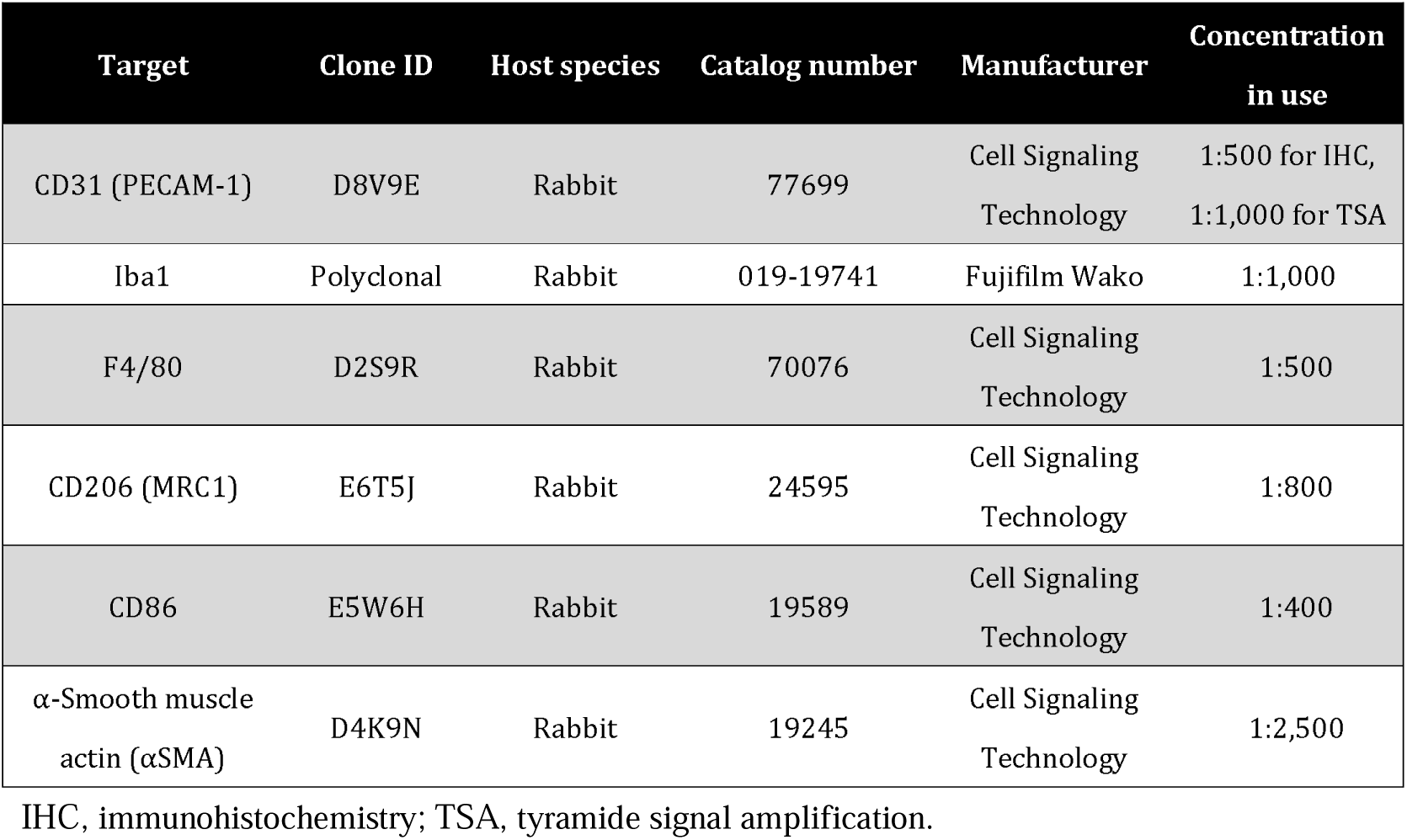
Antibody list.

| Target | Clone ID | Host species | Catalog number | Manufacturer | Concentration in use |
| --- | --- | --- | --- | --- | --- |
| CD31 (PECAM-1) | D8V9E | Rabbit | 77699 | Cell Signaling Technology | 1:500 for IHC, 1:1,000 for TSA |
| Iba1 | Polyclonal | Rabbit | 019-19741 | Fujifilm Wako | 1:1,000 |
| F4/80 | D2S9R | Rabbit | 70076 | Cell Signaling Technology | 1:500 |
| CD206 (MRC1) | E6T5J | Rabbit | 24595 | Cell Signaling Technology | 1:800 |
| CD86 | E5W6H | Rabbit | 19589 | Cell Signaling Technology | 1:400 |
| α-Smooth muscle actin (αSMA) | D4K9N | Rabbit | 19245 | Cell Signaling Technology | 1:2,500 |

IHC, immunohistochemistry; TSA, tyramide signal amplification.

Stained slides were scanned using a NanoZoomer S60 v2 slide scanner (Hamamatsu Photonics, Hamamatsu, Japan). Computational image analysis was performed using QuPath v0.6.0 [32]. Separate project files were generated for each staining marker and experimental group. Whole-slide images were imported as brightfield IHC/DAB images, and stain vectors were optimized for each staining set using the Estimate stain vectors function. Pixel classifiers were trained to segment viable tumor tissue, necrotic tissue, and background. Three slides per project were randomly selected and used as training samples for classifier generation. The trained classifiers were then applied to the remaining slides. Marker-specific thresholds were established to detect DAB-positive signals. Viable tumor areas were used to quantify DAB-positive signal area density.

Measured density values were analyzed using R v4.5.3 in RStudio v2026.01.2+418. PBS-treated and SQAP 4 mg/kg-treated tumors were compared within each HU-HSA-PDX model. Density values were log10-transformed before statistical testing. Two-sided Welch’s *t*-tests were used as the primary statistical test. *P*-values < 0.05 were considered statistically significant.

### CD31 and αSMA dual fluorescence analysis

Dual-fluorescence immunohistochemistry was performed on 3-μm sections obtained from the same formalin-fixed, paraffin-embedded tumor blocks used for H&E and conventional IHC analyses of HU-HSAPDX-3, HU-HSAPDX-6, and HU-HSAPDX-7. Tumors from the PBS and SQAP 4 mg/kg groups were analyzed. Sections were deparaffinized and subjected to heat-induced epitope retrieval in 1× ImmunoActive Antigen Retrieval Solution, pH 6.0 (Matsunami Glass Ind., Ltd.), using a 2100-Retriever (Aptum Biologics). Endogenous peroxidase activity was quenched with 0.3% hydrogen peroxide in methanol for 15 min at RT. Sections were then blocked with Blocking One Histo (Nacalai Tesque, Kyoto, Japan; Cat. No. 06349-64) for 25 min at RT. A sequential same-species tyramide signal amplification (TSA) workflow was used to detect CD31 and α-smooth muscle actin (αSMA) on the same section. CD31 was detected with an anti-CD31 (PECAM-1) rabbit monoclonal antibody (clone D8V9E; Cell Signaling Technology, Danvers, MA, USA; Cat. No. 77699), which detected murine endothelial cells but showed no cross-reactivity with canine HSA cells. αSMA was detected with an anti-αSMA rabbit monoclonal antibody (clone D4K9N; Cell Signaling Technology; Cat. No. 19245). Primary antibodies were diluted in SignalStain Antibody Diluent (Cell Signaling Technology; Cat. No. 8112) and incubated for 60 min at RT, followed by SignalStain Boost IHC Detection Reagent (HRP, Rabbit) (Cell Signaling Technology; Cat. No. 8114) for 30 min at RT. CD31 was labeled in the first round with Alexa Fluor 555 Tyramide (Thermo Fisher Scientific, Waltham, MA, USA; Cat. No. B40955). The antibody-HRP layer was then removed by heating the sections in fresh pH 6 retrieval buffer at 95-98 °C for 10 min, followed by cooling in the same buffer for 30 min. αSMA was labeled in the second round with Alexa Fluor 488 Tyramide (Thermo Fisher Scientific; Cat. No. B40953). Working tyramide solutions were prepared immediately before use in Tris-HCl buffer, pH 7.4, containing hydrogen peroxide.

Slides were mounted with ProLong Gold Antifade Reagent with DAPI (Cell Signaling Technology; Cat. No. 8961). A no-primary control and an inter-round carryover/strip control were included in each staining batch. Stained slides were scanned at 40× magnification (approximately 0.23 μm/pixel) using the NanoZoomer S60 v2 with DAPI, FITC, and TRITC/Cy3 filter channels. Multichannel images were analyzed in QuPath v0.7.0 with separate projects generated for each HU-HSAPDX model [32]. Viable tumor regions were manually annotated by reviewing DAPI nuclear patterns side by side with the corresponding H&E-stained section from the same tumor block. For each PDX model, a project-specific CD31 pixel classifier was trained using only the TRITC/Cy3 channel. The locked classifier was used both to measure CD31-positive area and to generate connected CD31-positive annotations with a minimum object area of 50 μm². Holes within these annotations were filled to create CD31_vessel_profile objects. A standardized vessel-wall band extending 3 μm inward and 3 μm outward from each CD31_vessel_profile boundary was then generated. All bands were merged into a non-overlapping union to prevent duplicate measurement of overlapping pixels. A second pixel classifier was trained using the DAPI, FITC, and TRITC/Cy3 channels to distinguish true αSMA-positive signal from erythrocyte autofluorescence and background. This classifier was applied to the non-overlapping wall-band union. To assess vascular maturity, αSMA coverage was calculated as the true αSMA-positive area divided by the evaluable wall-band area after excluding pixels classified as erythrocyte autofluorescence.

Results were analyzed using R v4.5.3 in RStudio v2026.01.2+418. CD31-positive area density was log10-transformed, and RBC-excluded αSMA coverage was logit-transformed before statistical testing. PBS- and SQAP 4 mg/kg-treated tumors were compared within each PDX model using two-sided Welch’s *t*-tests. Individual tumor values and group means ± standard error of the mean were plotted. *p-*values < 0.05 were considered statistically significant.

## Data availability statement

Raw sequencing data and processed expression matrices generated in this study have been deposited in the NCBI Gene Expression Omnibus under accession number GSE336925. Other data supporting the findings of this study are available from the corresponding author upon reasonable request.

## Acknowledgements

The authors thank Kenji Hosoya, Sangho Kim, Ryohei Kinoshita, and Ryo Owaki of the Veterinary Teaching Hospital, Hokkaido University, for providing the clinical tissue specimens used in this study. The authors are grateful to Dr. Hiroeki Sahara for his valuable advice on dual immunohistochemical staining to evaluate SQAP-associated changes in vascular maturity. The authors also gratefully acknowledge the canine patients with hemangiosarcoma whose tissue specimens were used in this study and their owners for consenting to the donation and use of these specimens for research. This work was supported by ASCO Co., Ltd.; a Japan Society for the Promotion of Science (JSPS) KAKENHI Grant-in-Aid for Scientific Research (B) (K.A.; 25K02164); and the Hokkaido University Frontier Foundation through public donations (K.A.).

## Author contributions

K.A. conceived the project, designed the experiments, performed most of the experiments, analyzed the data, and wrote the original draft. N.M., T.G., and K.H. performed histopathological and immunohistochemical analyses and contributed to data interpretation. All authors reviewed and edited the manuscript, approved the submitted version, and agreed to be accountable for the content of the work.

## Generative AI statement

The authors declared that Generative AI was used in the creation of this manuscript. OpenAI ChatGPT (model GPT-5.5 Pro; OpenAI) was used for English proofreading, manuscript organization, and methodological suggestions related to mRNA-seq, image-analysis, and statistical analyses. The tool was used only to support the authors’ writing and analytical workflow. AI-generated text, data, figures, or analysis results were not used verbatim. All outputs were critically reviewed, verified, revised, and finalized by the authors.

## References

1. Smith, A. N. Hemangiosarcoma in dogs and cats. Vet. Clin. North Am. Small Anim. Pract. 33, 533–552 (2003).

2. Kim, J. H., Graef, A. J., Dickerson, E. B. & Modiano, J. F. Pathobiology of hemangiosarcoma in dogs: Research advances and future perspectives. Vet. Sci. 2, 388–405 (2015).

3. Wendelburg, K. M. et al. Survival time of dogs with splenic hemangiosarcoma treated by splenectomy with or without adjuvant chemotherapy: 208 cases (2001–2012). J. Am. Vet. Med. Assoc. 247, 393–403 (2015).

4. Clifford, C. A., Mackin, A. J. & Henry, C. J. Treatment of canine hemangiosarcoma: 2000 and beyond. J. Vet. Intern. Med. 14, 479–485 (2000).

5. Batschinski, K., et al. Canine visceral hemangiosarcoma treated with surgery alone or surgery and doxorubicin: 37 cases (2005–2014). Can. Vet. J. 59, 967–972 (2018).

6. Tamburini, B. A. et al. Gene expression profiling identifies inflammation and angiogenesis as distinguishing features of canine hemangiosarcoma. BMC Cancer 10, 619 (2010).

7. Kim, J. H. et al. Interleukin-8 promotes canine hemangiosarcoma growth by regulating the tumor microenvironment. Exp. Cell Res. 323, 155–164 (2014).

8. Kim, J. H., et al. Hemangiosarcoma cells promote conserved host-derived hematopoietic expansion. Cancer Res. Commun. 4, 1467–1480 (2024).

9. Gulay, K. C. M. et al. Hemangiosarcoma cells induce M2 polarization and PD-L1 expression in macrophages. Sci. Rep. 12, 2124 (2022).

10. Kerboeuf, M. et al. Tumor-associated macrophages in canine visceral hemangiosarcoma. Vet. Pathol. 61, 32–45 (2024).

11. Choi, Y. & Jung, K. Normalization of the tumor microenvironment by harnessing vascular and immune modulation to achieve enhanced cancer therapy. Exp. Mol. Med. 55, 2308–2319 (2023).

12. Izaguirre-Carbonell, J., et al. Novel anticancer agent, SQAP, binds to focal adhesion kinase and modulates its activity. Sci. Rep. 5, 15136 (2015).

13. Iwamoto, H. et al. Inhibition of hypoxia-inducible factor via upregulation of von Hippel-Lindau protein induces ‘angiogenic switch off’ in a hepatoma mouse model. Mol. Ther. Oncolytics 2, 15020 (2015).

14. Sawada, Y. et al. Sulfoquinovosylacylpropanediol is a novel potent radiosensitizer in prostate cancer. Int. J. Urol. 22, 590–595 (2015).

15. Inamasu, E. et al. Anticancer agent α-sulfoquinovosyl-acylpropanediol enhances the radiosensitivity of human malignant mesothelioma in nude mouse models. J. Radiat. Res. 63, 19–29 (2022).

16. Takakusagi, Y. et al. A multimodal molecular imaging study evaluates pharmacological alteration of the tumor microenvironment to improve radiation response. Cancer Res. 78, 6828–6837 (2018).

17. Maeda, J., Sunada, S., Fukuhara, T. & Kato, T. A. A novel radiosensitizer α-sulfoquinovosyl-acylpropanediol (SQAP) inhibits DNA repair pathways and sensitize cells to cancer chemotherapeutic agents. Sci. Rep. 15, 41628 (2025).

18. Suzuki, T. et al. Lysine lactylation regulates ATF4-mediated stress responses under glucose starvation in canine hemangiosarcoma. Front. Vet. Sci. 13, 1734339 (2026).

19. Suzuki, T. et al. Lactate dehydrogenase is associated with cholesterol/lipid metabolism, and fluvastatin plus dipyridamole suppresses canine hemangiosarcoma growth in patient-derived xenograft models. Preprint at 10.64898/2026.03.03.709271 (2026).

20. Ruike, T. et al. Distribution and metabolism of ¹□C-sulfoquinovosylacylpropanediol (¹LC-SQAP) after a single intravenous administration in tumor-bearing mice. Xenobiotica 49, 346–362 (2019).

21. Percie du Sert, N., et al. The ARRIVE guidelines 2.0: updated guidelines for reporting animal research. PLoS Biol. 18, e3000410 (2020).

22. Dobin, A. et al. STAR: ultrafast universal RNA-seq aligner. Bioinformatics 29, 15–21 (2013).

23. Ahdesmäki, M. J., Gray, S. R., Johnson, J. H. & Lai, Z. Disambiguate: An open-source application for disambiguating two species in next generation sequencing data from grafted samples. F1000Research 5, 2741 (2017).

24. Li, H. et al. The sequence alignment/map format and SAMtools. Bioinformatics 25, 2078–2079 (2009).

25. Li, B. & Dewey, C. N. RSEM: accurate transcript quantification from RNA-Seq data with or without a reference genome. BMC Bioinformatics 12, 323 (2011).

26. Love, M. I., Huber, W. & Anders, S. Moderated estimation of fold change and dispersion for RNA-seq data with DESeq2. Genome Biol. 2014 15:12 15, 550 (2014).

27. Benjamini, Y. & Hochberg, Y. Controlling the false discovery rate: a practical and powerful approach to multiple testing. J. R. Stat. Soc. Ser. B Methodol. 57, 289–300 (1995).

28. Subramanian, A. et al. Gene set enrichment analysis: a knowledge-based approach for interpreting genome-wide expression profiles. Proc. Natl. Acad. Sci. USA 102, 15545–15550 (2005).

29. Mootha, V. K. et al. PGC-1α-responsive genes involved in oxidative phosphorylation are coordinately downregulated in human diabetes. Nat. Genet. 34, 267–273 (2003).

30. Liberzon, A. et al. Molecular signatures database (MSigDB) 3.0. Bioinformatics 27, 1739–1740 (2011).

31. Liberzon, A. et al. The molecular signatures database hallmark gene set collection. Cell Syst. 1, 417–425 (2015).

32. Bankhead, P., et al. QuPath: open source software for digital pathology image analysis. Sci. Rep. 7, 16878 (2017).

