## Supplemental Figures for "Sulfoquinovosylacylpropanediol monotherapy suppresses canine hemangiosarcoma patient-derived xenograft models with vascular remodeling"

### HU-HSAPDX-3

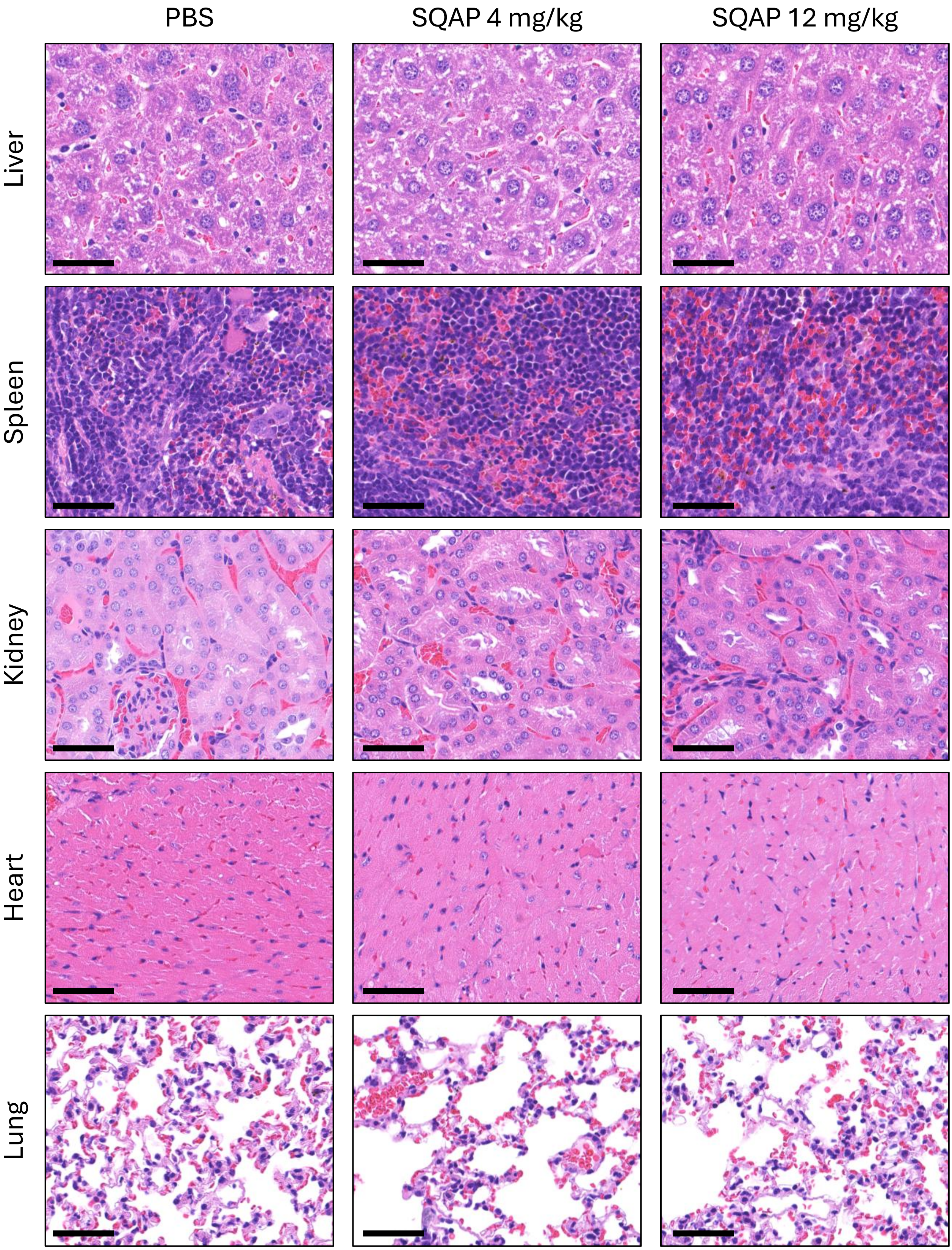


**Supplementary Figure S1. Histopathological evaluation of major organs from PBS/SQAP-treated HU-HSAPDX-3 mice.** Representative H&E-stained sections of the liver, spleen, kidney, heart, and lung from HU-HSAPDX-3 tumor-bearing mice treated with PBS, SQAP 4 mg/kg, or SQAP 12 mg/kg. Scale bars, 50 μm.

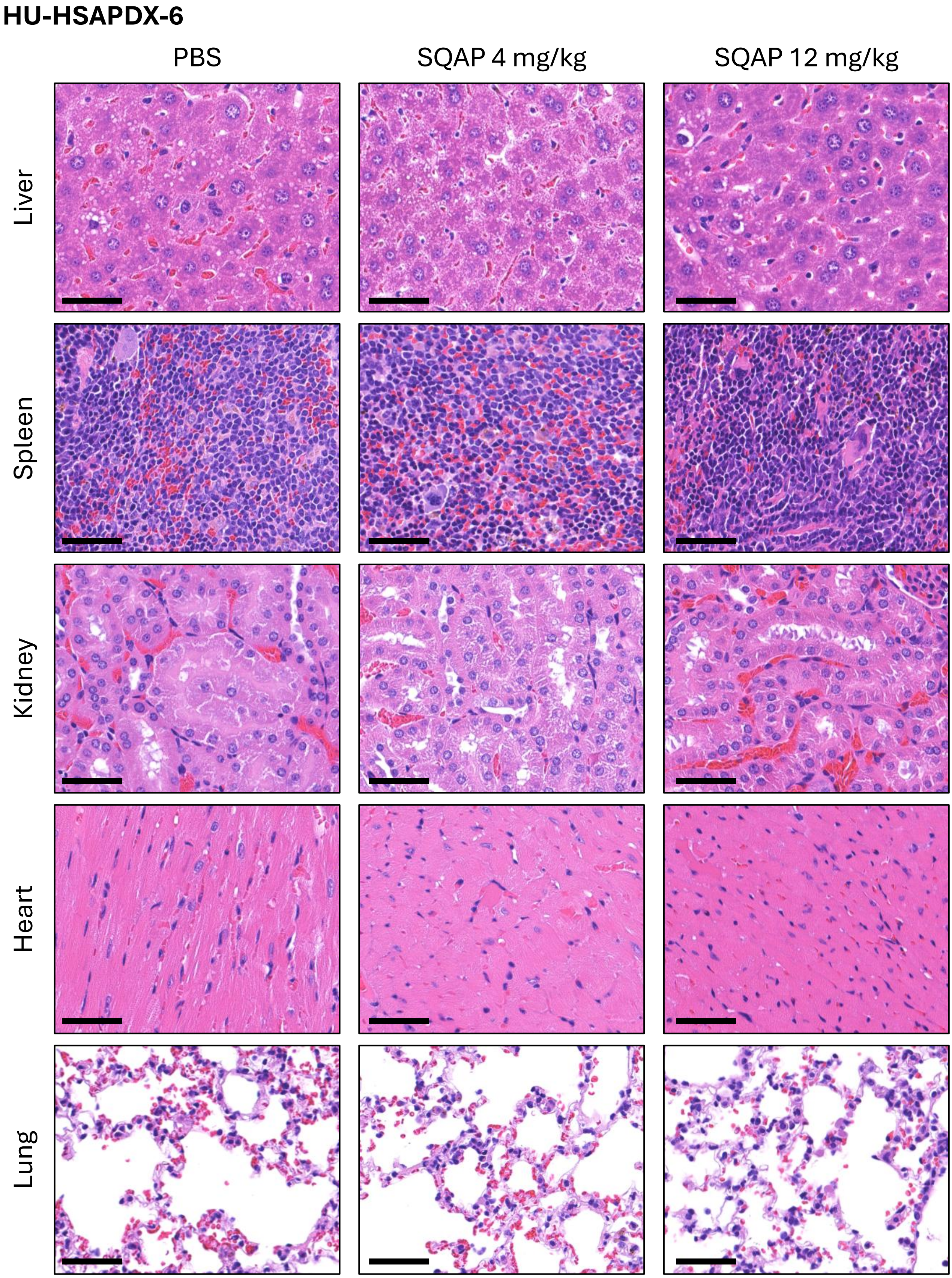


**Supplementary Figure S2. Histopathological evaluation of major organs from PBS/SQAP-treated HU-HSAPDX-6 mice.** Representative H&E-stained sections of the liver, spleen, kidney, heart, and lung from HU-HSAPDX-6 tumor-bearing mice treated with PBS, SQAP 4 mg/kg, or SQAP 12 mg/kg. Scale bars, 50 μm.

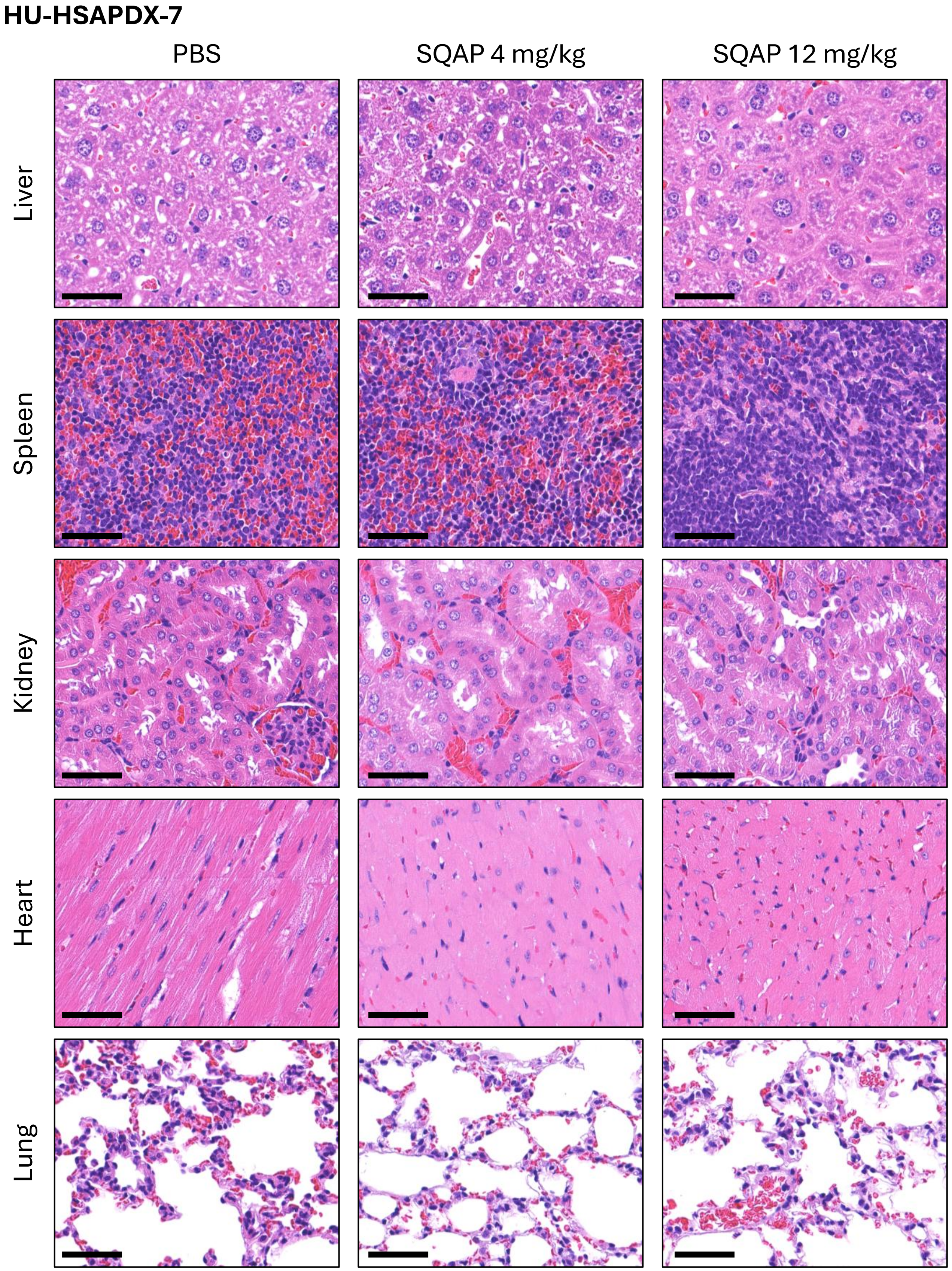


**Supplementary Figure S3. Histopathological evaluation of major organs from PBS/SQAP-treated HU-HSAPDX-7 mice.** Representative H&E-stained sections of the liver, spleen, kidney, heart, and lung from HU-HSAPDX-7 tumor-bearing mice treated with PBS, SQAP 4 mg/kg, or SQAP 12 mg/kg. Scale bars, 50 μm.
